# The planning horizon for movement sequences

**DOI:** 10.1101/2020.07.15.204529

**Authors:** Giacomo Ariani, Neda Kordjazi, J. Andrew Pruszynski, Jörn Diedrichsen

## Abstract

When performing a long chain of actions in rapid sequence, future movements need to be planned concurrently with ongoing action. However, how far ahead we plan, and whether this ability improves with practice, is currently unknown. Here we designed an experiment in which healthy volunteers produced sequences of 14 finger presses quickly and accurately on a keyboard in response to numerical stimuli. On every trial, participants were only shown a fixed number of stimuli ahead of the current keypress. The size of this viewing window varied between 1 (next digit revealed with the pressing of the current key) and 14 (full view of the sequence). Participants practiced the task for five days and their performance was continuously assessed on random sequences. Our results indicate that participants used the available visual information to plan multiple actions into the future, but that the planning horizon was limited: receiving information about more than 3 movements ahead did not result in faster sequence production. Over the course of practice, we found larger performance improvements for larger viewing windows and an expansion of the planning horizon. These findings suggest that the ability to plan future responses during ongoing movement constitutes an important aspect of skillful movement. Based on the results, we propose a framework to investigate the neuronal processes underlying simultaneous planning and execution.

**Significance Statement:** Although skill learning has typically focused on the training of specific movement sequences, practice improves performance even for random sequences. Here we hypothesize that a fundamental aspect of skilled sequential behavior is the ability to plan multiple actions into the future, both before and during execution. By controlling the amount of visual information available for motor planning, we show that people plan at least three movements beyond current action and that this planning horizon expands with practice. Our findings suggest that coordinating ongoing movement and planning of future actions is an essential component of skilled sequential behavior and offer testable predictions for the neural implementation of online motor planning.

## Introduction

Humans exhibit a wide range of behaviors, from whole-body activities like running or riding a bike, to fine dexterous skills like writing or typing on a keyboard. Many of such skills share one common feature: they are comprised of a series of separate motor elements that are strung together in quick succession to form longer and more complex sequences of movements (Lashley, 1951). When learning a new sequential skill, people usually need many hours of practice to achieve fluidity in performance (Ericsson et al., 1993). With practice, sequence production becomes quicker, more accurate, and less effortful (Diedrichsen and Kornysheva, 2015; Krakauer et al., 2019; Rhodes et al., 2004; Verwey, 1994), leading in the long run to the skillful behaviors typically observed in elite athletes (Yarrow et al., 2009).

Previous studies of motor sequence learning have largely focused on the training of specific movement sequences (Cohen et al., 1990; Kornysheva et al., 2019, 2013; Mantziara et al., 2020; Verwey, 2001; Verwey et al., 2014; Verwey and Abrahamse, 2012; Willingham, 1999; Aaron L. Wong et al., 2015). However, many sequences we execute in everyday life are not fully predictable and practice improves performance even for random or untrained sequential movements (Ariani et al., 2020; Waters-Metenier et al., 2014; Wiestler et al., 2014). Some of these sequence-general improvements arise because participants learn to translate individual visual stimuli into motor responses and to execute these responses more quickly (Ariani and Diedrichsen, 2019; Hardwick et al., 2019). Such improvements in single responses benefit the production of all sequences, including random ones.

In the present study, we focus on a second core ability that benefits the production of unpredictable sequences: the ability to plan future movements ahead of time. Planning of movements before their initiation, here referred to as *preplanning*, has been studied extensively (Churchland et al., 2010; Cisek and Kalaska, 2010; Haith et al., 2016; Kaufman et al., 2014; Rosenbaum, 1980; Rosenbaum et al., 2007, 1987; Aaron L Wong et al., 2015). However, long or complex movement sequences are unlikely to be fully preplanned, so planning of the remaining elements must continue throughout sequence production – a process that we have recently named *online planning* (Ariani and Diedrichsen, 2019). Take the example of a basketball player dribbling up the court. The player needs to control a continuous flow of movements (e.g., to keep the dribble alive) while scouting the court and planning future movements depending on the actions of both teammates and opposing defenders. Some evidence for online planning has been observed for a range of behaviors, such as reading (Rayner, 2014, 1998; Rayner and Reingold, 2015), walking (Matthis and Fajen, 2014), sequential reaching (Säfström et al., 2014, 2013) and path tracking (Bashford et al., 2018). However, to what extent the motor system plans upcoming movements during sequence production (i.e., the horizon of online planning) remains poorly understood.

Here we asked 1) how far the benefit of planning ahead extends beyond current execution, and 2) whether this planning horizon can be improved with practice. To answer these questions, we used a discrete sequence production (DSP) task, in which participants performed random sequences of 14 keypresses with their right hand in response to numerical cues. We manipulated how many digits participants could see ahead of the current keypress. Viewing window size ranged from 1 (only the next movement is cued, as in the serial reaction time task, SRTT) to 14 (the entire sequence shown at once, as in the DSP task). Participants practiced producing varying sequences over 5 days. This design allowed us to examine the horizon of both pre- and online planning in sequence production, as well as the influence of practice on the planning horizon.

## Methods

### Participants

Seventeen right-handed neurologically healthy volunteers (8 women, 9 men; age 18–36 years, mean 25.81 years, SD 5.09 years) were recruited for this study. Handedness was assessed with the Edinburgh Handedness Inventory (mean 82.81, SD 18.07). Individuals participated in 5 sessions of practice (2 hours each, on 5 separate days). All participants provided written informed consent and were naive to the purposes of the study. Two participants abandoned the study after the first session of practice. One participant had an unusually high error rate (> 30%, while every other participant managed to keep the error rate < 20%, as per instructions). These 3 participants were excluded from successive analyses (final N = 14). For one of the remaining 14 participants, age and handedness data was missing, and for another participant eye tracking data was missing. All experimental methods were approved by the Research Ethics Board at Western University.

### Apparatus

Participants placed their right hand on a custom-made keyboard (Fig. 1A), with a force transducer (Honeywell FS series) mounted underneath each key. The keys were immobile and measured isometric finger force production. The dynamic range of the force transducers was 0-16 N and the resolution 0.02 N. A finger press/release was detected when the force crossed a threshold of 1 N. The forces measured from the keyboard were low pass filtered, amplified, and sent to PC for online task control and data recording. Additionally, we recorded monocular left eye movements using an SR Research EyeLink 1000 desk-mounted eye tracker. Eye movements were recorded at a rate of 500 Hz. Participants sat approximately 40 cm away from a 20” (50.8 cm) screen. Numerical stimuli were shown in white against a black background, horizontally aligned in a single line, and spanned 13.5cm for an entire sequence (∼19° of visual angle). Individual digits were about 1 cm tall, 0.5 cm wide, and spaced 1 cm apart (from center to center, ∼1.43° of visual angle).

**Figure 1.**
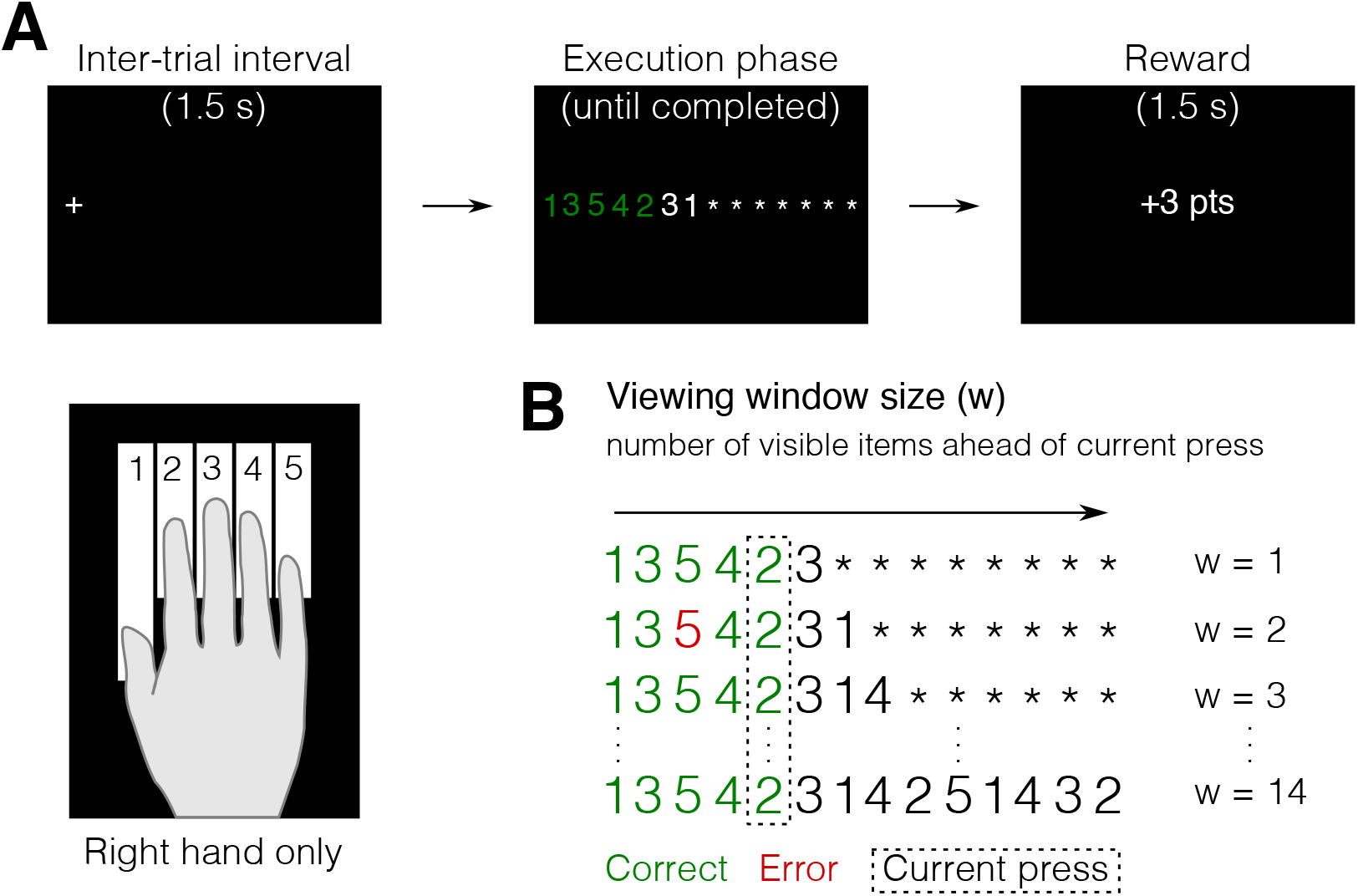
Varying viewing window in a discrete sequence production (DSP) task. **A.** Example trial in a DSP task with viewing 2 items ahead of the current keypress, while the remaining items are masked by asterisks. **B.** Viewing window size (w) manipulation, from w = 1 (equivalent to a simple reaction time task), to w = 14 (display of the entire sequence at once). The arrow indicates the from-left-to-right direction of response order. Participants could start each sequence whenever they felt ready and were rewarded on the basis of their movement time (MT, the time from the first keypress to the release of the last key in the sequence).

### Procedure

In each of the five practice sessions, participants sat in front of a computer screen with their right hand on the keyboard, and their chin placed on the eye tracker chinrest. The task required participants to produce sequences of keypresses in response to numerical cues appearing on the screen (numbers 1 to 5, corresponding to fingers of their right hand, thumb to little finger, respectively) as quickly and accurately as possible (Fig. 1A). On every trial, only a fixed number of digits (viewing window size, w) were revealed to the participants, while the rest were masked with asterisks (Fig. 1B). The masked digits were revealed to the participant as they proceeded, from left to right, with the presses in each sequence. The window size varied within the domain of w = {1, 2, 3, 4, 5, 6, 7, 8, 14}, and was randomized across trials within every block. As an attentional pre-cue, at the beginning and at the end of every trial, during the inter-trial interval (ITI, 1.5 seconds), participants were presented with a fixation cross on the location of the first digit in the sequence. This fixation was used to account for possible drifts in trial-by-trial calibration of eye-to-digit location mapping. Within every block of trials, participants were instructed to keep their chin on the eye tracker chinrest at all times and to minimize head movements. With every press, subjects received feedback about the correctness of their action: the white numbers turned either green or red and were accompanied by either a high-pitch or a low-pitch sound for correct and incorrect presses, respectively.

To motivate participants to improve in the task, they were rewarded with points based on their performance after each trial. Points were awarded on the basis of sequence movement time (MT) and execution accuracy. MT corresponded to the time interval between making the first press in the sequence to releasing the last press in the sequence. Accuracy was calculated as 1 – error rate (proportion of error trials in a block) in percentage. Specifically, a trial was considered an error if it contained one or more incorrect presses, for which participants received 0 points. Correct sequences were rewarded with at least 1 point. Finally, participants were awarded 3 points if 1) a sequence was correct and 2) MT 5% or more faster than a specific time threshold. This time threshold was designed to get increasingly difficult adjusting to every subject’s speed throughout training. It would decrease by 5% from one block to the next if two performance criteria were met: median MT in the current block faster than best median MT recorded hitherto, and mean error rate in the last block ≤ 15%. If either one of these criteria was not met, the thresholds remained unchanged. At the end of each block, participants received feedback on their error rate, median sequence MT, total points obtained during the block, and total points obtained during the session. Subjects were asked to try to maintain an error rate below 15%.

In the original design, we intended to compare also how the ability to plan ahead might affect partially familiar (structured) sequences. Therefore, each one of the 5 practice sessions consisted of 8 blocks (27 trials each) of 14-item sequences and 3 blocks (60 trials each) of specific short 3-/4-item segments that composed the structured sequences. One-third of the trials in the sequence blocks were randomly generated by random shuffles of the digits 1 to 5. The remaining two-thirds of the trials were structured sequences. As the results from the structured sequences turned out to be hard to interpret, the present paper focuses only on the completely unfamiliar, random sequences.

### Control experiment to measure the limits imposed by visual acuity and crowding effects

To determine how far out from the fovea the digit stimuli could be recognized, we ran a short control experiment. In a modified version of the task, we asked an independent sample of participants (N = 3) to fixate on the first digit of a random series of 14 digits and report all digits with finger presses, while maintaining fixation on the first digit. The task was non-speeded, as we wanted to determine how many digits could be recognized, given human limits in visual acuity (Rayner, 1975) and the limits imposed by attentional crowding effects (Levi, 2008). Participants were instructed to press the fingers corresponding to the digits they could see, and guess when they were not sure anymore. Each participant repeated this task for 6 blocks with 27 trials each. We calculated the probability of making the correct keypress as a function of distance from the fixated digit. Chance performance was *p* = 0.2 (1 out of 5 fingers).

### Data analysis

Data were analyzed with custom code written in Matlab (The MathWorks, Inc., Natick, MA). To evaluate the speed of sequence production, we inspected the time intervals between different keypresses. Reaction times (RT) were defined as the time from stimulus onset to first press (i.e., the first crossing of the 1 N force threshold). Note that participants were not instructed to react particularly fast. Instead, they could take as much time as they wanted until they felt ready to start. MTs were defined as the time between the first press and the release of the last press in the sequence (i.e., the time between the first and the last crossing of the force threshold). Finally, we calculated inter-press intervals (IPI) between subsequent pairs of presses in the sequence (i.e., the time interval between every two consecutive crossings of the force threshold). Unless otherwise noted, we used within-subject repeated measures ANOVAs and 2-sided paired samples *t-*tests for statistical inference in assessing the effects of viewing window or practice on RT, MT, and IPI. Error trials were excluded from data analysis. To provide meaningful error bars for within-subject comparison, the standard error for each condition was calculated on the residuals after subtracting the mean across conditions for each participant. This way, the error bars visualized the size of the relevant error term in a repeated-measures ANOVA.

To describe the relationship between MT and the viewing window size, we used the following exponential model:

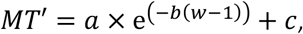

where *MT’* is the predicted MT for a given viewing window size w. Note that for w = 1, the function reduces to the initial value of the exponential, *MT’* = *a* + *c*. The asymptote is given by *c* and the slope by *b*. This model was then fit to the MT data of each participant using Matlab’s `nlinfit()` function, which implements the Levenberg-Marquardt nonlinear least-squares algorithm. We determined the effective planning horizon (*w\**), by finding the window size for which the predicted MT of the participant had dropped 99% of the difference between w = 1 and the asymptote, i.e., by solving the equation for w:

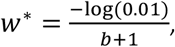

where the 0.01 arises from the criterion of the 99% drop (i.e., 1% above the *MT’* asymptote). The improvement in effective planning horizon with practice was then assessed by fitting the model to the data of each participant on each day and comparing *w\** between day 1 and day 5 with a within-subject 2-sided paired samples *t-*test. While the use of a 99% criterion is arbitrary, changes in this criterion only scale the effect planning horizon by a specific value but do not change the outcome of the statistical analysis.

### Analysis of eye movements

To assess changes in fixation strategies, we estimated the eye position with respect to each finger press in the sequence as follows. For each trial in each block, we mapped the calibrated eye position (in eye tracker units) to the digits on the screen to calculate the digit (*D*_*t*_) on which the eye was currently fixated at the time of each keypress:

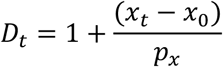

Where ***x***_***t***_ is the eye’s current horizontal position at the time of press (median position within a 25 ms time window around the keypress) in eye tracker units, ***x***_*0*_ is the median horizontal position of the eye at the beginning of the trial (i.e., the position of the fixation cross / first digit), and ***p***_***x***_ is a normalizing unit constant throughout a block of trials, used to convert eye tracker units to digit positions. Finally, we computed the estimated eye position at the time of each keypress by subtracting the position corresponding to the keypress (1 to 14) from ***D***_***t***_ and plotted this estimate against each keypress position in the sequence (Fig. 7). In other words, a value of 0 means that the eye was exactly on the digit at was currently pressed, and a value of +1 would indicate that the eyes were a full digit ahead the currently pressed digit.

## Results

### Preplanning of future movements speeds up sequence production

First, we assessed the benefit of being able to plan future finger movements on sequence production. To determine this, we varied the amount of available information and tested how this window size affected the speed of performance. On average, across all days of practice, larger window sizes produced shorter MTs (Fig. 2A), as confirmed by the highly significant main effect of window size on MT in a repeated measures ANOVA (*F*_8,104_ = 176.980, *p* < 10e-10). This finding suggests that the availability of visual information allows for preplanning of sequential actions into the future, which in turn reduces MT. Interestingly, this benefit appeared to plateau around a window size of 3 or 4. Indeed, when we compared the MT of each viewing window to the average MT for larger window sizes, we found a significant difference for w = 3 vs w > 3 (*t*_13_ = 4.644, *p* = 4.591e-04), but not for w = 4 vs w > 4 (*t*_13_ = 2.083, *p* = 0.058). To obtain an individual measure of the planning horizon, we fit an exponential model to the MT curve of each participant (Fig. 2B, see Methods).

**Figure 2.**
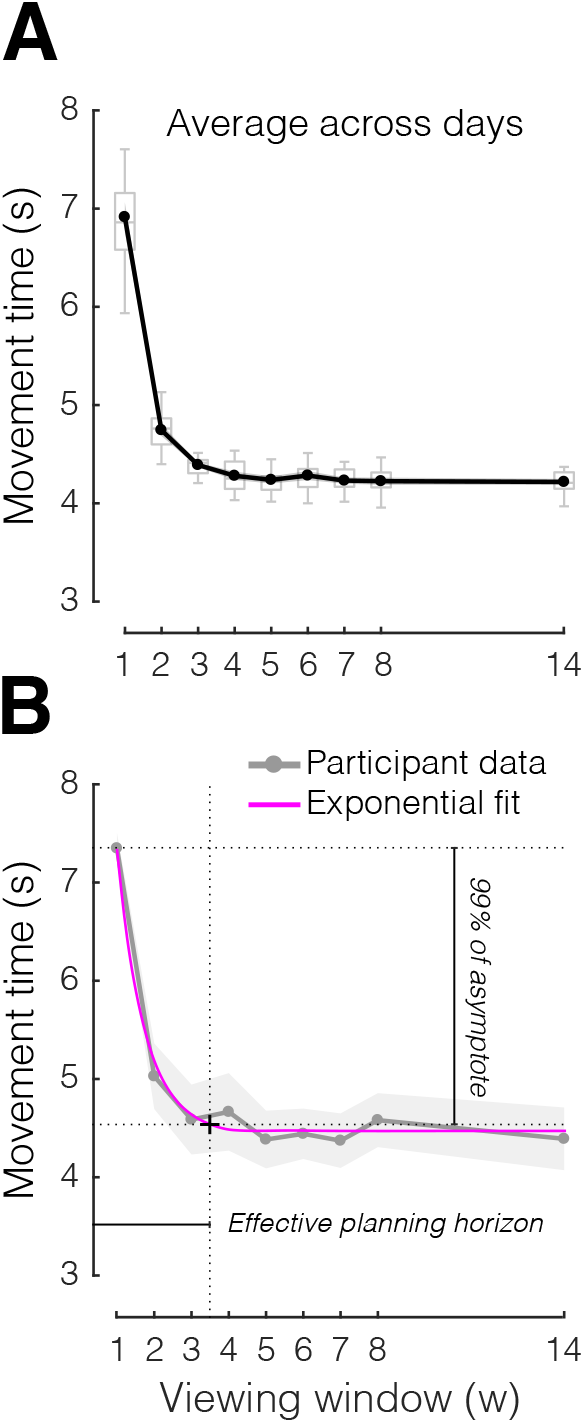
The benefit of planning ahead on sequential performance. **A.** Average movement time as a function of viewing window, across the 5 days of practice. **B.** Method used to estimate the effective planning horizon. Example data from one participant (gray) is fit to an exponential model (magenta). The intersection between performance at 99% of asymptote and the exponential fit was chosen as criterion to determine the effective planning horizon. Box plots show the median and whole range of individual data points (one per participant). Shaded areas reflect standard error of the mean.

Next, we set an arbitrary criterion on the exponential (99% of the MT drop to the asymptote) to establish the individual effective horizon of each participant. This analysis revealed a mean effective planning horizon of 3.58 ± 0.28 items ahead of the current item, indicating that, on average, participants were able to plan at least 3 keypresses into the future.

Was this limitation in planning horizon simply due to a perceptual limitation? Obviously, the drop-off of visual acuity from the fovea to the periphery (Rayner, 1975) could limit the ability of the participants to identify more visually presented letters simultaneously. Moreover, even if acuity is sufficient, the presentation of multiple digits can lead to “crowding”, an ubiquitous attentional effect that impairs the ability to recognize visual objects in clutter (Levi, 2008). To test both of these possible limitations for our display, we conducted a control experiment (see Methods) and found that participants were able to accurately identify up to 6 items to the right of fixation (∼6 cm) with more than 95% accuracy (mean accuracy for 7^th^ item = 97.67% ± 0.78 95% CI). Thus, given that eye position was on average only slightly to the right of the currently pressed digit (see analysis of eye movement strategies below), it seems very unlikely that peripheral visual acuity and crowding can explain the limitation in planning horizon.

### Practice expands the planning horizon

We then asked whether practicing sequences would affect the ability to plan future movements by comparing performance at the beginning (day 1) and at the end (day 5) of practice (Fig. 3A). We observed that MT improved across all window sizes (main effect of day: *F*_1,13_ = 18.004, *p* = 0.001). Significant improvements were found even for a window size of 1 (MT difference day 1 vs. day 5: 586 ± 262 ms; *t*_13_ = 2.234, *p* = 0.022). This condition was, in essence, a serial reaction time task, where each cue was only presented after the preceding key was pressed, meaning that participants were forced to serially cycle through the planning and execution of every press, with no possibility for planning ahead. Therefore, MT improvements for a window size of 1 must be a consequence of 1) better stimulus identification, 2) better stimulus-response (S-R) mapping, or 3) better execution (i.e., motor implementation) of single responses.

**Figure 3.**
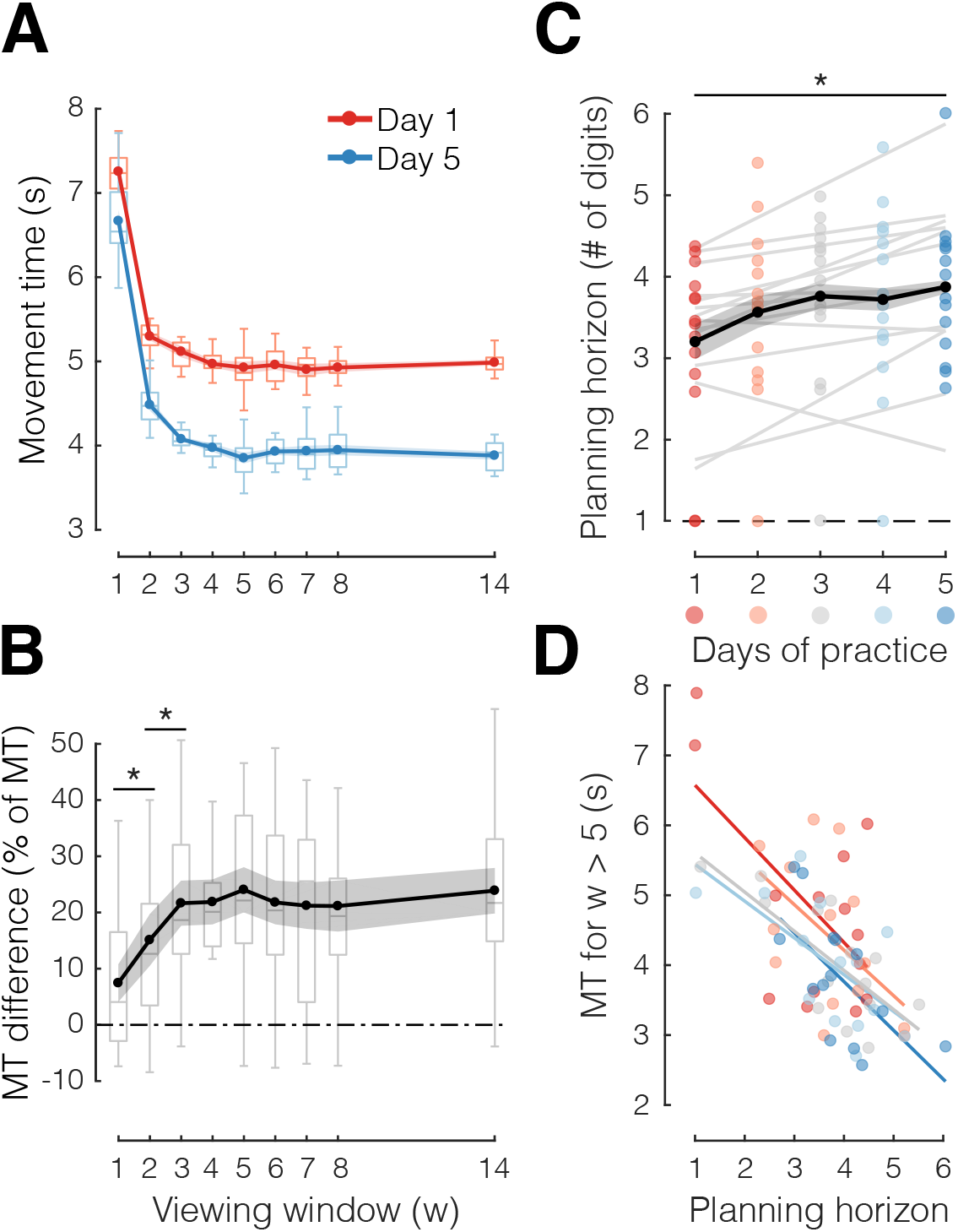
The effective planning horizon increases with practice. **A.** Average movement time (MT) as a function of viewing window (w), separately for early (day 1, red) and late (day 5, blue) stages of sequence practice. **B.** Difference in performance (sequence MT) between early and late practice (data in A), normalized by average MT for each w, as a function of w. **C.** Mean effective planning horizon (estimated as shown in Fig. 2B) for each day of practice.**D.** Correlation between mean MT for w > 5 (which enabled planning ahead) and mean effective planning horizon, separately for different days (days 1 to 5 in gradient red to blue). For this analysis, planning horizon was estimated on odd blocks and MT on even blocks. Dots reflect individual data points (one per participant). Box plots show the median and quartiles of group data. Shaded areas reflect standard error of the mean. *p < 0.05, two-tailed paired-samples t-tests.

If learning was restricted to improvement in any of these three processes, we would predict equal MT improvement across all window sizes, given that stimulus identification, S-R mapping, and execution are necessary steps across all viewing windows. Contrary to prediction, we found a significant interaction between window size and stage of practice (day 1 vs. day 5; *F*_8,104_ = 3.220, *p* = 0.003). Furthermore, when we directly inspected the MT improvement (percentage change relative to average MT for each w, Fig. 3B), we found significantly larger gains for larger viewing windows (w = 2 vs. w = 1: *t*_13_ = 3.338, *p* = 0.005; w = 3 vs. w = 2: *t*_13_ = 2.722, *p* = 0.017), until again the gains plateau for w = 4 or larger (w = 4 vs. w = 3: *t*_13_ = 0.113, *p* = 0.912). Thus, although responses to single items improved with practice, this improvement cannot explain why performance benefits were more pronounced for larger window sizes. Instead, the additional performance benefit must arise because participants became more efficient at using the advance information provided by larger viewing windows.

Part of this increased efficiency may be due to expansion of the planning horizon (i.e., how far ahead participants were able to plan). Indeed, we also found evidence that with practice, participants planned further into the future. When we determined the effective planning horizon for each participant and day (Fig. 2B) using an exponential fit (see Methods), we found that the planning horizon expanded from 3.20 to 3.88 digits ahead of current action between day 1 and day 5 (paired-samples *t*-test, *t*_13_ = 2.840, *p* = 0.014, Fig. 3C). Despite the high inter-subject variability, out of 14 participants, only 2 show a negative slope when regressing the effective planning horizon onto the day of practice. Furthermore, there was no relationship between the average planning horizon and the slope (i.e., the rate of change) across days (r = 0.28, *p* = 0.3376), suggesting that participants with small and large horizons improved similarly.

While significant, the increase of planning horizon by less than a digit appears to be quite small. How important is a larger planning horizon for faster performance with practice? To obtain insight into this question, we looked at the inter-subject variability within each training day: We correlated our measure of planning horizon for each participant with their mean movement time for larger viewing windows (w > 5) for each day of practice. To avoid any statistical dependency between these parameters induced by measurement noise, the effective planning horizon was estimated on data from odd blocks and the mean movement time was estimated on data from even blocks. This analysis (Fig. 3D) revealed a clear negative correlation between the two measures for each day of practice (all r < - 0.54, all *p* < 0.05). Roughly speaking, from this analysis, we would expect that the observed increase in effective planning horizon by 0.68 items should lead, on average, to a decrease in movement time of about 435 ms, which corresponds to ∼82% of the improvement from day 1 to day 5 for w = 14 (∼1110 ms) that was not explained by improvements in single responses (i.e., ∼580 ms for w = 1). Thus, participants became better at planning future movements in advance, and this improvement could potentially be explained by the increased ability to plan more actions into the future.

Note that faster MTs for larger window sizes did not occur at the expense of reduced accuracy in performance. On average, the percent accuracy of presses remained roughly constant around 85-90% across all viewing window conditions. We found no significant main effect of window size (*F*_8,104_ = 1.182, *p* = 0.317), practice stage (*F*_1,13_ = 0.325, *p* = 0.578), or interaction between the two factors (*F*_8,104_ = 0.548, *p* = 0.818).

Taken together, these results show that participants became faster in sequence production by getting better at 1) making single responses (involving stimulus identification, S-R mapping, or execution) and 2) exploiting available information to plan more upcoming movements in advance.

### Reaction times increase with the amount of preplanning

If participants invested time in preplanning the first few elements of each sequence, then we would expect this to be reflected in their reaction times: namely, participants should start a sequence earlier when presented with a smaller window size, and later for larger window sizes, since they would be preparing more of the upcoming keypresses. Even though fast RTs were not required by the task, participants likely tried to balance the benefit of getting more points with the benefit of finishing the experiment more quickly. On average, across all days (Fig. 4A), larger viewing windows resulted in slower RTs (*F*_8,104_ = 4.563, *p* = 8.726e-05).

**Figure 4.**
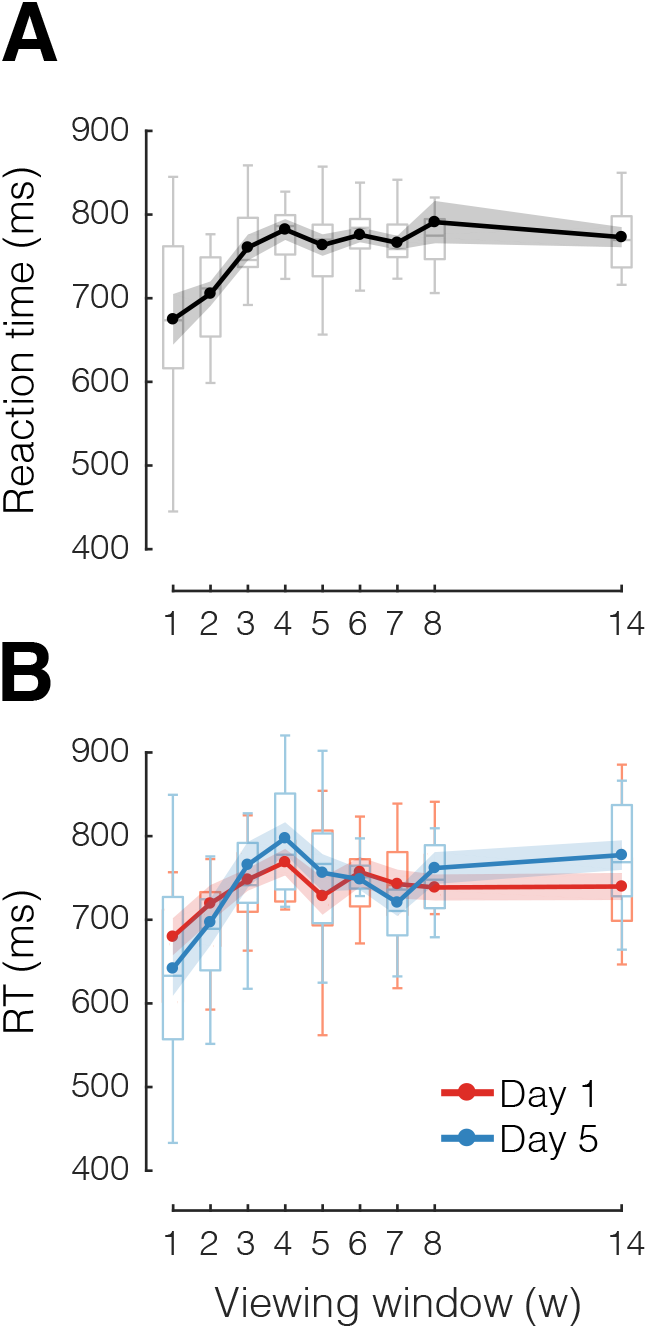
Longer reaction times for larger viewing windows. **A.** Average reaction time as a function of viewing window. **B.** Subset of data in A, separating between early (day 1, red) and late (day 5, blue) stages of practice. Box plots show the median and whole range of individual data points (one per participant). Shaded areas reflect standard error of the mean.

However, as observed for MTs, RTs appeared to plateau for window sizes larger than 3. Thus, even though participants could see more than 3 elements on the screen and had virtually unlimited time to preplan, they initiated the sequence in approximately 700-800 ms from cue onset.

Did this relationship between RTs and the amount of available information change with practice? When we compared RTs across early and late stages of practice (Fig. 4B) we found no indication that, late in practice, participants waited longer to initiate a sequence (*F*_1,13_ = 0.012, *p* = 0.913), or that their strategy changed over time (no interaction between practice stage and window size: *F*_8,104_ = 1.187, *p* = 0.314).

### Planning ahead continues during sequence production

So far, our results have indicated that participants improve their ability to perform random sequences of finger movements by becoming more efficient in using the information provided by larger window sizes. However, it remains unclear whether participants got better at planning movements before sequence production (preplanning), during sequence production (online planning), or both. To distinguish the contributions of preplanning and online planning to performance improvements, we examined the time intervals between individual presses in a sequence (i.e., the IPIs). The rationale behind this analysis is that short IPIs reflect an increased readiness to press (i.e., better planning) than long IPIs. If all keypresses were equally well prepared (e.g., as in the case of w = 1, which does not allow participants to plan ahead), then all IPIs within a sequence should roughly have the same duration depending on the serial RT (null hypothesis, Fig. 5A). Alternatively, if only early presses in a sequence can be fully preplanned, then only these IPIs should be significantly shorter, and later IPIs revert to serial RT speed (Fig. 5B). Finally, if online planning continues in parallel with execution, we should expect an effect of window size also on mid to late IPIs (Fig. 5C).

**Figure 5.**
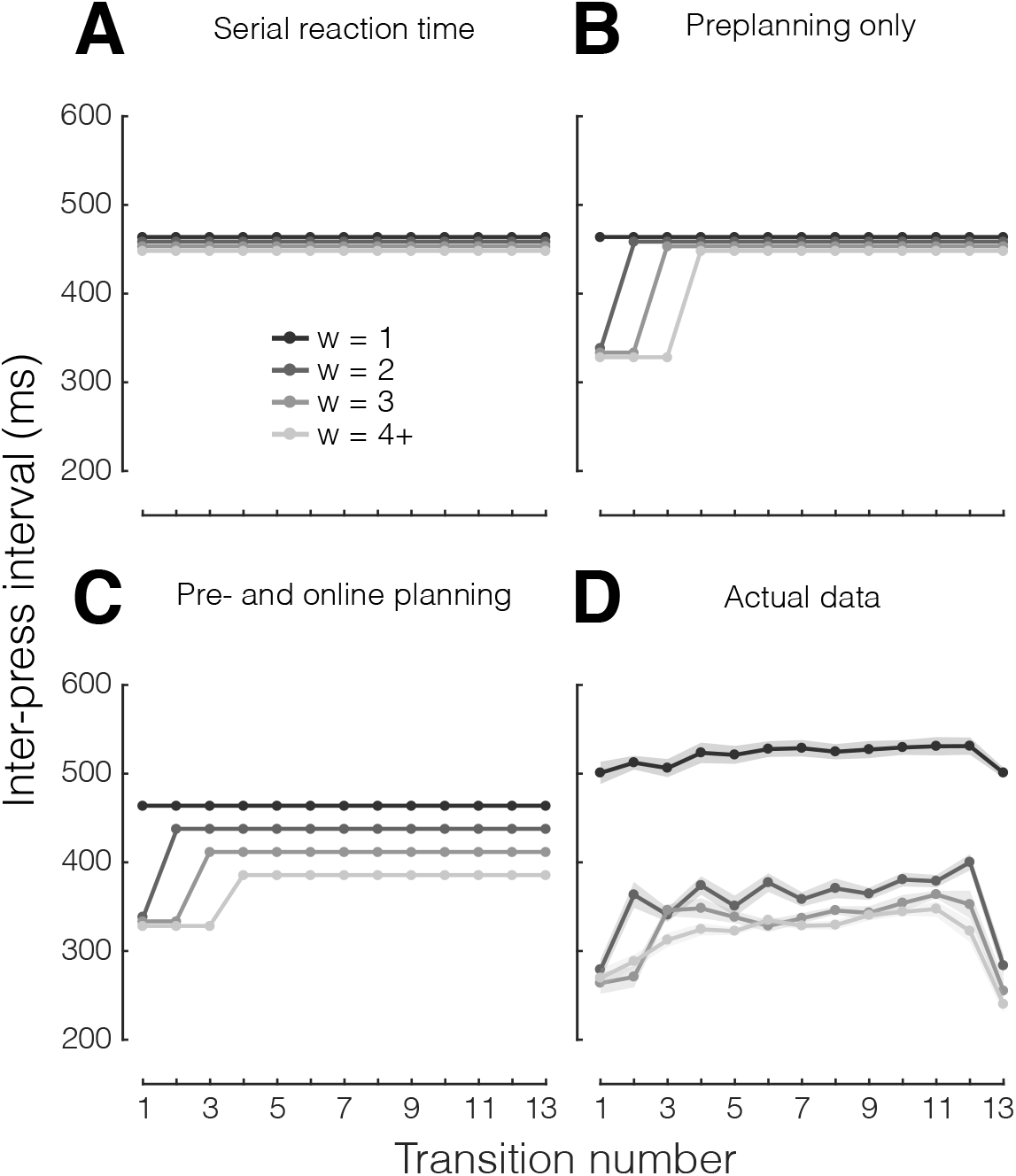
Predictions and analysis of inter-press intervals (IPIs). Average inter-press interval (IPI) as a function of transition number within each sequence, separately for viewing window size (w, different shades of gray). 4+ indicates w ≥ 4. **A.** Prediction 1 (null hypothesis): no effect of w, all IPIs roughly in the same range. **B.** Prediction 2: Fast early IPIs reflect the benefit of preplanning, but for unplanned keypresses the benefit of viewing ahead is minimal. **C.** Prediction 3: even mid to late IPIs benefit from larger w, indicating that both pre- and online planning are contributing to fast sequence production. **D.** Actual group data of mean IPIs for each keypress transition, separately for each viewing window.

In light of these predictions, we first inspected the IPIs averaging across practice stages (Fig. 5D). For a window size of 1, all IPIs had approximately the same duration (∼500 ms), reinforcing the idea that for w = 1, each keypress is selected, planned, and executed independently. In contrast, for window sizes larger than 1, we found a clear effect of IPI placement (i.e., finger transition number within the sequence) on IPI duration (*F*_12,156_ = 33.111, *p* < 10e-10). Specifically, the first and last IPIs were consistently performed much faster than the middle IPIs, regardless of the size of the viewing window (W > 1). For w = 2, the first IPI (first 2 finger presses) was faster than subsequent IPIs; for w = 3, the first two IPIs (first 3 finger presses) were faster than subsequent IPIs. For w > 3, this preplanning advantage appeared to be spread over the first 3 finger transitions. This pattern of results indicates that the initial speed up reflects the fact the visible digits can be preplanned during the reaction time, and hence are executed faster.

Consistent with RT and MT data, preplanning does not seem to improve further beyond a window size of 3. This observation reinforces the idea that participants preplanned at least the first three movements of each finger sequence. Once all preplanned keypresses are executed, planning must continue online, slowing down later IPIs. Thus, the slower IPIs in the middle of the sequence mostly reflect limits in the speed of online planning. When we restricted our analysis to these middle IPIs (transitions 5 to 12), the differences between w = 1 and w = 2 (*t*_13_ = 19.557, *p* = 5.037e-11), between w = 2 and w = 3 (*t*_13_ = 5.013, *p* = 2.374e-04), and between w = 3 and w = 4 remained significant (*t*_13_ = 2.182, *p* = 0.048). This indicates that, just like preplanning, online planning benefits from having visual information about up to 3 presses into the future, thus highlighting clear parallels between the two processes.

We also observed that, consistent across all window sizes greater than 1, the last IPI was executed much more quickly than preceding IPIs. Currently, we do not have a definitive answer about the reasons for this result. One idea is that participants tend to select and plan the last 2 presses as a unit. These presses can then be executed very quickly, as no more movements need to be planned after those two (which frees up planning capacity). Alternatively, participants could optimize the last two presses from an execution biomechanics perspective. Given that no subsequent movements are needed, participants do not have to maintain a specific hand posture that would be required for fast execution of successive movements. Instead, they are free to optimize their hand posture for comfort and speed only in regard to making the last two presses.

### Both pre- and online planning improve with practice

Finally, we asked whether practice effects on MT are more likely related to improvements in preplanning, online planning, or both. From day 1 to day 5 (Fig. 6A), we observed significant main effects of practice stage on IPI duration on both early (IPI 1-3: *F*_1,13_ = 17.623, *p* = 0.001) and middle IPIs (IPI 5-12: *F*_1,13_ = 15.988, *p* = 0.002). To quantify the relative contributions of preplanning and online planning, we carried a separate analysis (Fig. 6B) averaging across IPIs that were more likely preplanned (IPI 1 for w = 2, IPI 1-2 for w = 3, and IPI 1-3 for w ≥ 4), or not (the remaining IPIs for each viewing window condition, which had to be planned online). For w = 1, only the first press, but not the first IPI, can be preplanned. Therefore, we cannot attribute any of the observed improvements to either sequence preplanning or online planning. Instead, eventual improvements need to arise from improved visual identification, S-R mapping, or execution. We computed the IPI difference between day 1 and day 5 for these three categories, normalized it by the average IPI duration across days (separately for each category), and plotted it against viewing window size (Fig. 6B). This analysis confirmed that IPIs became faster with practice even for w = 1 (one-sample *t-*test vs zero difference: *t*_13_ = 2.305, *p* = 0.038). Additionally, we found clear further improvements in IPI duration for w > 1: compared to w = 1, these effects were present both for the IPIs that were likely preplanned (*t*_13_ = 4.028, *p* = 0.001), and for those that relied on online planning (*t*_13_ = 6.009, *p* = 4.379e-05). There was no significant difference between preplanning and online planning in terms of learning improvements (*F*_1,13_ = 1.141, *p* = 0.305), nor was there an interaction between planning process and viewing window (*F*_2,26_ = 1.000, *p* = 0.382). Thus, preplanning and online planning appear to have similar capacity limits and to benefit similarly from practice in sequence production. Moreover, given that on day 5 participants did not spend more time preplanning than they did on day 1 (Fig. 4B), improvements in IPI with practice (Fig. 6A) reinforce the idea that participants could make use of more visual information in roughly the same amount of preparation time (i.e., with comparable RTs).

**Figure 6.**
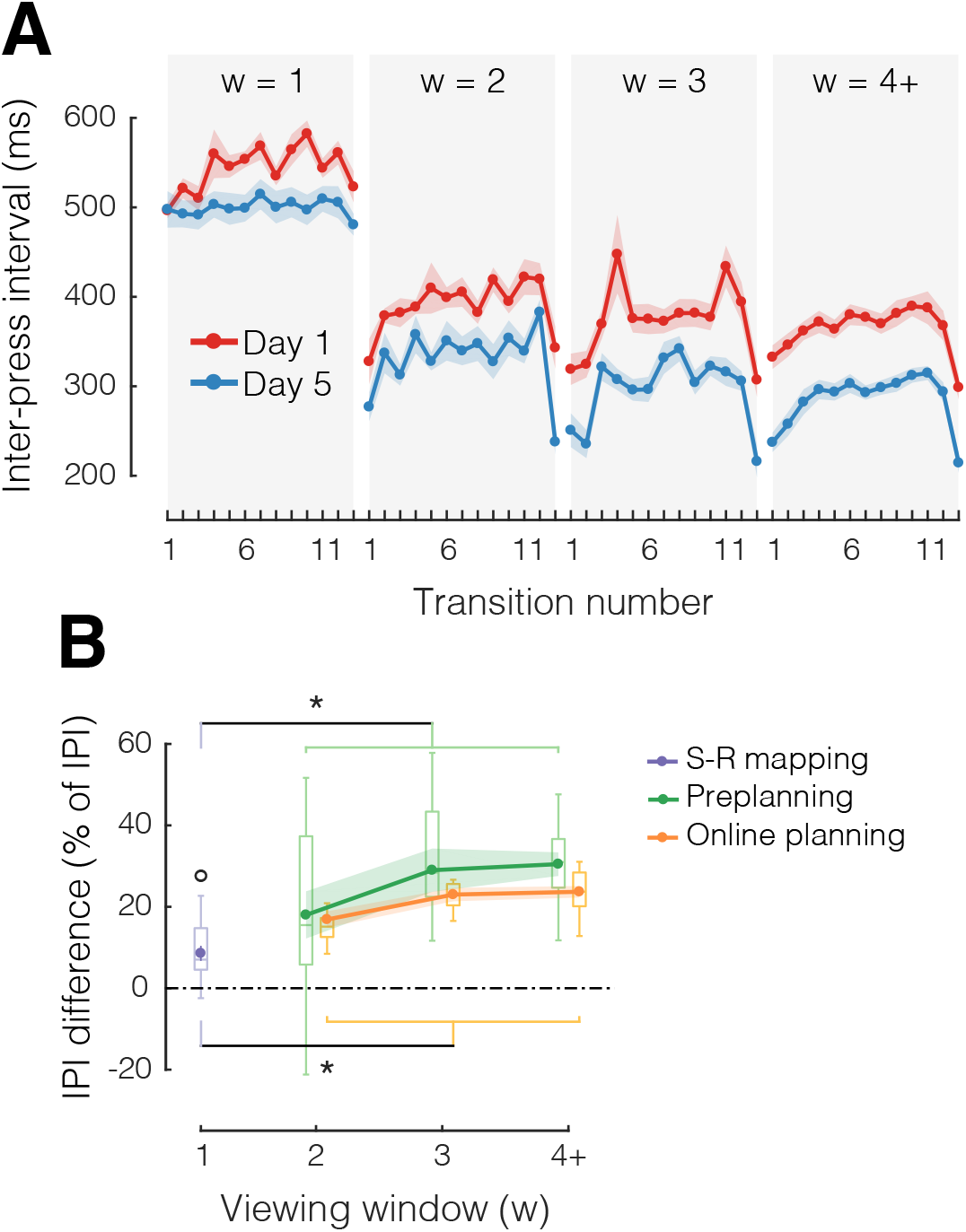
Improvements in pre- and online planning with practice. **A.** Mean IPI as a function of transition number, separately for practice stage (day 1, red; day 5, blue) and viewing window size (w = 1, 2, 3, 4+ in separate plots). **B.** Average IPI difference between day 1 and day 5, normalized by average IPI for each day, separately for each w and planning process (S-R mapping, purple; Preplanning, green; Online planning, orange). Box plots show the median and whole range of individual data points (one per participant). Shaded areas reflect standard error of the mean. *p < 0.05, two-tailed paired-samples t-tests; °p < 0.05, two-tailed one sample t-test.

### Increases in planning horizon are not explained by changes in eye movement strategies

To determine to what degree changes in fixation strategies could cause improvements in the planning horizon, we analyzed eye tracking data. To assess potential learning effects in fixation strategies, we estimated eye position at the time of each keypress in the sequence (see Methods). We then plotted this estimated eye position at the time of each press relative to the digit that was produced (Fig. 7). A value of 0 in means that the eye was exactly on the digit at was currently pressed, and a value of +1 would indicate that the eyes were a full digit ahead the currently pressed digit. For statistical comparison across days, we selected the keypresses in the middle of a sequence (presses 3-10) given that those are most likely to be influenced by fixation strategies. A 2-by-4 repeated measures ANOVA revealed a significant main effect of viewing window size (*F*_3,36_ = 17.815, *p* < 0.0001), indicating that subjects tended to look further ahead when more information was available. However, no main effect of day (*F*_1,12_ = 0.624, *p* = 0.445) or interaction between day and window size was found (*F*_3,36_ = 2.739, *p* = 0.058), indicating that fixation strategies were not the cause of improvements in the effective planning horizon. The results were similar even when including all of the keypresses (1-14): significant main effect of window size (*F*_3,36_ = 10.691, *p* < 0.0001), but no main effect of day (*F*_1,12_ = 0.730, *p* = 0.410), or interaction between the two (*F*_3,36_ = 1.779, *p* = 0.169). Together with our data showing that perceptual limitations cannot account for the limited planning horizon (see above), these results argue that participants increased their ability to preplan more actions by overcoming a central (cognitive-motor) bottleneck, rather than a purely perceptual bottleneck.

**Figure 7.**
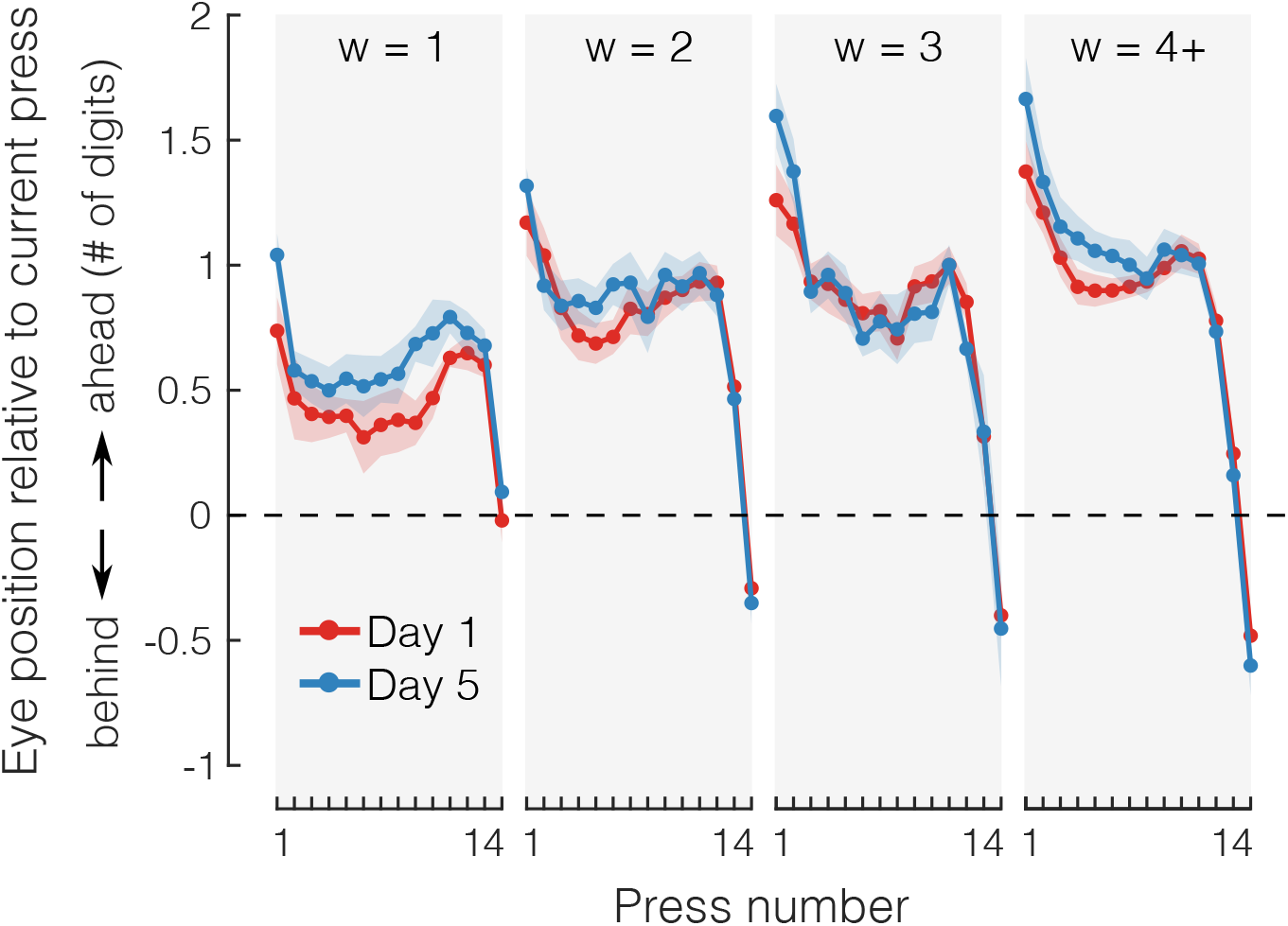
Eye movement strategies do not change with practice. Mean eye position relative to current press at the time of press as a function of press number, for different days of practice (Day 1, red; Day 5, blue) and for different viewing windows (w, gray shaded areas). Viewing windows of size of 4 or larger were grouped together as 4+. Shaded areas reflect standard error of the mean. Dashed line indicates eye position that would be equivalent to looking directly at the digit corresponding to the current finger press. Positive numbers indicate that the eyes are further ahead than the current press, and negative numbers indicate that the eyes are lagging behind.

## Discussion

The ability to move and simultaneously plan future movements is a fundamental yet underappreciated faculty of the human brain. By manipulating the amount of visual information available for motor planning in a discrete sequence production task, we show that participants planned multiple actions (at least 3) into the future (Fig. 2-4). Practice led to larger gains in speed for larger window sizes (MT difference, Fig. 3B), as well as increases in the horizon of sequence planning (exponential fit, Fig. 3C). In-depth analysis of the inter-press intervals (Fig. 5-6) revealed that enhanced planning of future actions was present both before (preplanning) and during (online planning) sequence production.

### Fast sequence production relies on the speed of online planning

Before voluntary movements can be performed, they need to be planned (Bock and Arnold, 1992; Cisek and Kalaska, 2004, 2002; Crammond and Kalaska, 2000, 1994; Keele, 1968; Keele and Summers, 1976; Kerr, 1978; Rosenbaum, 1980), at least to some degree (Ames et al., 2014; Cisek and Kalaska, 2010). However, many real-life motor skills require quick sequences of movements that are not always predictable. Proficiency in such skills depends on our ability to select and plan future movements both before and during sequence production. To investigate this ability, we used a viewing window paradigm that varied the amount of information available for planning the next movements. We replicate previous findings of anticipatory planning in the context of random sequences (Herbort and Butz, 2009; Rhodes et al., 2004; Rosenbaum et al., 1987), and longer reaction times when more information for planning is available (Henry and Rogers, 1960). Critically, once such preplanning reaches capacity, the execution of later elements in the sequence slows down, which we interpret as evidence that successive movements need to be planned online. By varying the available time for preplanning, in a previous paper (Ariani and Diedrichsen, 2019) we showed that this was the case even for relatively short (e.g., 5-item) and well-known (e.g., trained) sequences – only the first 3 elements were fully planned prior to execution. Further evidence for online planning comes from a wide range of activities in which visual information is used for planning. For example, when participants make anticipatory eye movements to future targets in reading (Rayner, 1978), sequential reaching (Säfström et al., 2014), and object manipulation (Johansson et al., 2001). More directly, a recent unpublished study tackled the horizon of online planning by restricting the viewing window in a continuous manual tracking task (Bashford et al., 2018). Together, these studies support our view that the ability of the motor system to deal with a stream of incoming stimuli while producing motor responses (i.e., online planning) enables skillful performance of movement sequences.

### Motor planning has a limited capacity

We found that the span of the planning horizon (∼3-4 movements) was smaller than the typical amount of information that can be stored in short-term memory (Cowan, 2010; Miller, 1956). However, more recent theories of short-term memory posit that capacity is not limited by a fixed number of items, but rather by finite attentional resources that can be flexibly allocated across multiple items (Bays and Husain, 2008; Luck and Vogel, 1997). Similarly, characterizing the planning horizon as having a hard, discrete limit (i.e., a specific number of movements) may not be the best description. A more realistic model would assert that cognitive resources of a central bottleneck (Pashler, 1994) are mainly allocated to the next upcoming movement, with a decaying distribution for elements further into the future (Fig. 8A). Such soft distribution of central resources to multiple tasks is consistent with classical models of the Psychological Refractory Period (McLeod, 1977; Smith, 1967; Welford, 1952). In the context of sequence preplanning, this idea is also consistent with the Competitive Queueing (CQ) hypothesis (Averbeck et al., 2006, 2002; Kornysheva et al., 2019; Mantziara et al., 2020; Rhodes et al., 2004): the first element would be preplanned the most, with subsequent elements being prepared to a decreasing degree. Eventually, subjects run out of resources and start executing responses. Completing preceding movements frees up new resources that can be allocated to plan successive movements online. The discrepancy between the short-term memory span and the planning horizon may reflect the fact that planning a movement takes up more central resources than remembering a digit.

**Figure 8.**
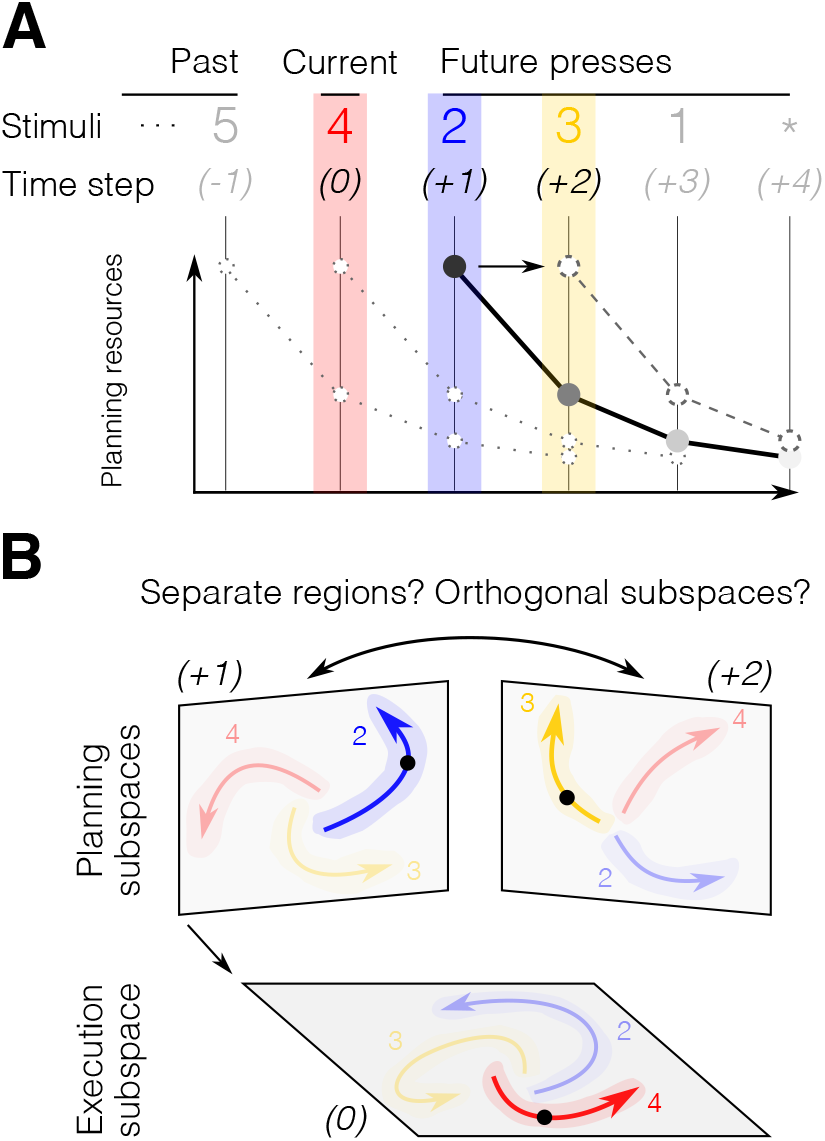
Planning capacity and hypothetical neural implementation of sequential behavior. **A.** The “soft” horizon of sequence planning depends on the amount of resource available. In this illustrative example, most resources are invested in the planning of the immediately upcoming press (+1, 2, blue). The further in time a press is from the current press (0, 4, red), the smaller the corresponding planning investment. Once a press has been initiated, the resources are redistributed by shifting the planning curve one step ahead, thus allowing for continuous online planning of future presses (e.g., +2, 3, yellow). **B.** Hypothetical neuronal population activity in brain regions involved in planning and execution processes. Each plane refers to an independent subspace of the multidimensional population activity with possible neural trajectories for the current action (0), and future actions (+1, +2), color-coded as in A. Shaded areas reflect single trial variability. The current neural state is indicated by a black dot. Planning of the next (+1) and future actions (+2) may evolve in separate regions or in orthogonal subspaces within the same region.

Importantly, the limits in planning horizon appear to be of central (cognitive) origin, rather than strictly perceptual or motor. As shown by the first (and last) IPI in a sequence, participants are in principle capable of executing key presses faster than the asymptote reached even for large window sizes. Additionally, perceptually, participants are able to identify 6 digits to the right of their eye fixation, which is at least a full digit more than their average planning horizon.

### The horizon of motor planning expands with practice

If the capacity of motor planning depends on a soft, flexible horizon, we can ask whether this limit can be improved with practice. In agreement with a previous study (Bashford et al., 2018), we found that practice had expanded the span of the planning horizon. Our conclusion was based on two key observations: 1) the benefit of seeing further ahead was greater later in practice (significant interaction between w and day on MT); 2) the influence of window size on MT can be described with an exponential function whose decay rate decreased with practice (change in the slope of the exponential). Speed improvements that are independent of the amount of available information can be attributed to improved stimulus identification, S-R mapping, or implementation of single responses (Ariani and Diedrichsen, 2019; Haith et al., 2016; Hardwick et al., 2019). As participants become more fluent at translating numbers on the screen into finger movements, each individual press is executed more quickly, thus contributing to faster sequence production across all window sizes. The greater performance benefits for larger window sizes together with the expansion of the effective planning horizon indicate that participants improved their ability to make better use of advance information. Given the nature of the exponential fit, it is hard to separate the relative contributions of planning efficiency (i.e., benefitting more from a specific window size) and larger horizons to such improvements. Unlike previous studies that examined sequence-specific effects in sequence production (Ariani and Diedrichsen, 2019; Berlot et al., 2020; Verwey, 2001; Verwey and Wright, 2004; Wiestler and Diedrichsen, 2013), here we focused on random sequences. Note that, because of this, the observed practice effects cannot be explained by the formation of specific chunking structures previously proposed as a way to deal with the complexity of planning long movement sequences (Popp et al., 2020; Ramkumar et al., 2016). Instead, we found that even without prior experience with a specific sequence, people can improve in the motor planning processes that underlie sequence production. In other words, practice effects are not only about learning *what* sequence to produce, but also about learning *how* to coordinate execution and planning efficiently.

### Implications for the neural control of sequential movements

The present study provides behavioral evidence that online planning constitutes a central component of motor skill. More speculatively, it raises the question of how planning and execution processes can simultaneously occur in the brain without interfering with each other. One possibility is that planning and execution processes take place in separate but communicating anatomical areas (Fig. 8B), such as the dorsal premotor cortex (PMd) for planning and the primary motor cortex (M1) for execution. However, several studies have reported signals related to movement planning also in brain structures responsible for movement (Ames et al., 2019, 2014; Ariani et al., 2018; Crammond and Kalaska, 2000; Elsayed et al., 2016), with signals often mixed even within single neurons (Alexander and Crutcher, 1990; Evarts and Tanji, 1976; Prut and Fetz, 1999; Riehle and Requin, 1989). Therefore, a more likely scenario is that planning and execution occur in overlapping neuronal populations, but occupy orthogonal subspaces of the multidimensional neuronal code (Fig. 8B), such that planning activity does not trigger motor output (Kaufman et al., 2014; Lara et al., 2018; Zimnik and Churchland, 2020). Finally, an intriguing possibility that should be investigated in future studies is that multiple future movements could be planned as an integrated packet (i.e., as a movement chunk), such that there is only one, shared, planning subspace, with specialized code for specific transitions (i.e., 2-3) in movement sequences. Namely, for random sequences, and provided a large enough viewing window, one could imagine that participants plan ahead until they can see and execute that chunk before moving on to the planning of what has since been revealed in a discontinuous fashion (which is revealed by longer and non-homogeneous inter-press intervals). Likewise, in the context of known (i.e., learned) or predictable sequences, it is possible that participants could recall and anticipate an upcoming chunk of information by only seeing part of it, thus planning and executing that chunk before moving on to the next. Some evidence compatible with such an integrated code has been reported in studies of the SMA (Hoshi and Tanji, 2004; Tanji and Shima, 1994), which found neurons sensitive to specific sequences of actions. However, most neurophysiological studies have focused on planning-related signals before movement onset (preplanning). Therefore, it remains an open question how the neural substrates for planning change when the same neuronal population has to concurrently control an ongoing movement (i.e., online planning). Our behavioral results highlight notable similarities between preplanning and online planning: both processes led to faster performance when participants had a chance to plan up to 3 upcoming sequence elements, with diminishing gains for larger window sizes. Additionally, practice-related improvements were comparable between early IPIs (mostly preplanning) and late IPIs (online planning). These similarities suggest that preplanning and online planning may rely on the same neural process (i.e., motor planning) happening in different contexts, either in isolation before movement initiation or in parallel with execution.

## Acknowledgements

The authors thank Eva Berlot for comments on earlier versions of the manuscript.

## Notes

**Funding sources**. This work was supported by a James S. McDonnell Foundation Scholar award and a NSERC Discovery Grant (RGPIN-2016-04890) to J.D., and the Canada First Research Excellence Fund (BrainsCAN). J.A.P is funded by the Canada Research Chairs program.

### Competing Interest Statement

The authors have declared no competing interest.

